# A refined set of rRNA-targeted oligonucleotide probes for *in situ* detection and quantification of ammonia-oxidizing bacteria

**DOI:** 10.1101/2020.05.27.119446

**Authors:** Michael Lukumbuzya, Jannie Munk Kristensen, Katharina Kitzinger, Andreas Pommerening-Röser, Per Halkjær Nielsen, Michael Wagner, Holger Daims, Petra Pjevac

**Affiliations:** University of Vienna, Centre for Microbiology and Environmental Systems Science, Division of Microbial Ecology, Vienna, Austria; Center for Microbial Communities, Department of Chemistry and Bioscience, Aalborg University, Aalborg, Denmark; Max Planck Institute for Marine Microbiology, Bremen, Germany; University of Hamburg, Institute of Plant Science and Microbiology, Microbiology and Biotechnology, Hamburg, Germany; Joint Microbiome Facility of the Medical University of Vienna and the University of Vienna, Vienna, Austria; University of Vienna, The Comammox Research Platform, Vienna, Austria

**Keywords:** Fluorescence *in situ* hybridization, ammonia-oxidizing bacteria, nitrification, wastewater treatment plants, oligonucleotide probes

## Abstract

Ammonia-oxidizing bacteria (AOB) of the betaproteobacterial genera *Nitrosomonas* and *Nitrosospira* are key nitrifying microorganisms in many natural and engineered ecosystems. Since many AOB remain uncultured, fluorescence *in situ* hybridization (FISH) with rRNA-targeted oligonucleotide probes has been one of the most widely used approaches to study the community composition, abundance, and other features of AOB directly in environmental samples. However, the established and widely used AOB-specific 16S rRNA-targeted FISH probes were designed up to two decades ago, based on much smaller rRNA gene sequence datasets than available today. Several of these probes cover their target AOB lineages incompletely and suffer from a weak target specificity, which causes cross-hybridization of probes that should detect different AOB lineages. Here, a set of new highly specific 16S rRNA-targeted oligonucleotide probes was developed and experimentally evaluated that complements the existing probes and enables the specific detection and differentiation of the known, major phylogenetic clusters of betaproteobacterial AOB. The new probes were successfully applied to visualize and quantify AOB in activated sludge and biofilm samples from seven pilot- and full-scale wastewater treatment systems. Based on its improved target group coverage and specificity, the refined probe set will facilitate future *in situ* analyses of AOB.

## 1. Introduction

Nitrification, a key process in the biogeochemical nitrogen cycle, is the microbially mediated oxidation of ammonia to nitrite and subsequently to nitrate. For many decades nitrification was perceived as a process always performed by two functional groups of aerobic, chemolithoautotrophic microorganisms in cooperation: the ammonia oxidizing bacteria (AOB) and archaea (AOA), which oxidize ammonia to nitrite, and the nitrite oxidizing bacteria (NOB) that oxidize nitrite to nitrate (Bock and Wagner, 2001; Daims et al., 2016; Könneke et al., 2005). Only recently, complete ammonia oxidizers (comammox organisms) have been discovered, which carry out the entire nitrification process alone (Daims et al., 2015; van Kessel et al., 2015).

Currently, all known canonical AOB, oxidizing ammonia to nitrite, belong to one of two phylogenetic lineages within the Proteobacteria. Gammaproteobacterial AOB include the genus *Nitrosococcus*, which is halophilic and occurs in marine habitats and salt lakes (Campbell et al., 2011), and the genus *Nitrosoglobus*, which contains acidotolerant AOB from acidic soils (Hayatsu et al., 2017). In contrast, the AOB that usually dominate in terrestrial and freshwater ecosystems belong to the family *Nitrosomonadaceae* within the (now obsolete, Parks et al., 2018) taxonomic class of Betaproteobacteria (Prosser et al., 2014). Here, we refer to these organisms as β-AOB. All cultivated and described members of the family *Nitrosomonadaceae* are chemolithoautotrophic AOB from the genera *Nitrosomonas* and *Nitrosospira* (Prosser et al., 2014). Representatives of β-AOB are found in almost all oxic environments but are particularly successful in nutrient rich habitats such as fertilized soils or eutrophic freshwater sediments, and also the majority of wastewater treatment plants (WWTPs) (Bollmann et al., 2014; Di et al., 2010; Fan et al., 2011; Verhamme et al., 2011). The low abundance of AOA in activated sludge from most municipal WWTPs (Gao et al., 2013; Mußmann et al., 2011; Wells et al., 2009) has recently been attributed to their sensitivity to copper limitation caused by chemical complexation of copper by organic compounds (Gwak et al., 2019). This effect likely contributes to the commonly observed predominance of β-AOB in these engineered environments.

Despite their ubiquity, β-AOB have proven to be exceptionally fastidious and recalcitrant to cultivation. Hence, the number of β-AOB isolates with standing in nomenclature (Parte, 2014) remains low (n=14) despite their broad environmental distribution and high ecological significance. To overcome this problem, researchers use cultivation-independent molecular techniques for studying various aspects of AOB diversity and ecophysiology. One of the most commonly applied molecular methods is rRNA-targeted fluorescence *in situ* hybridization (FISH). This approach uses rRNA-targeted oligonucleotide probes, which are covalently linked to fluorescent dyes and hybridize to the ribosomal RNA of specific microbial populations (Amann et al., 1995; DeLong et al., 1989). The resulting fluorescence signal allows the *in situ* detection and visualization of target organisms in environmental samples. FISH has numerous applications in microbial ecology, which include the *in situ* abundance quantification of populations (Daims and Wagner, 2007; Wagner et al., 1994), and quantitative analyses of the spatial distribution of microorganisms in biofilms and other structurally complex samples (Almstrand et al., 2013; Daims et al., 2006; Dolinšek et al., 2013; Schillinger et al., 2012; Welch et al., 2016). Combinations of FISH with chemical imaging techniques like microautoradiography (Lee et al., 1999), Raman microspectroscopy (Fernando et al., 2019; Huang et al., 2007), and nanometer scale secondary ion mass spectroscopy (NanoSIMS) (Berry et al., 2013; Musat et al., 2012) even permit cultivation-independent physiological studies of discrete microbial populations. FISH can also be used together with bioorthogonal noncanonical amino acid tagging (BONCAT), which is another powerful approach to detect metabolically active microorganisms *in situ* (Hatzenpichler et al., 2014).

FISH has been applied since 1995 (Wagner et al., 1995) in numerous studies to investigate β-AOB in aquatic systems, especially WWTPs, and has proven to be of immense value in this context. For example, *Nitrosomonas* (formerly “*Nitrosococcus*”) *mobilis* was identified as the dominant AOB in an industrial WWTP (Juretschko et al., 1998) and later isolated from activated sludge by FISH-assisted screening and propagation of sorted microcolonies (Fujitani et al., 2015). FISH and image analysis were used to quantify the spatial localization patterns of β-AOB in nitrifying biofilms (Almstrand et al., 2013; Gruber-Dorninger et al., 2015, p.; Maixner et al., 2006). By combining FISH detection with microsensor measurements of substrate concentration gradients, both the distribution and activities of β-AOB in biofilms were studied (Okabe et al., 1999; Schramm et al., 1998). Application of this approach to calculate volumetric reaction rates even revealed the *in situ* whole-cell kinetics of uncultured β-AOB (*Nitrosospira* spp.) (Schramm et al., 1999). In another study, FISH and quantitative PCR (qPCR) were used to detect β-AOB in granular activated sludge. These data were the basis for two ecophysiological models, which address the observed (and unexpected) higher *in situ* abundances of NOB over AOB in the granules (Winkler et al., 2012). FISH was also used to quantify the abundance dynamics of diverse β-AOB and anaerobic ammonium oxidizers (anammox) in ammonium- or urea-fed enrichments, revealing different substrate preferences of the populations (Sliekers et al., 2004), and to analyze the spatial organization of β-AOB and anammox organisms in nitrogen-removing biofilms (Pynaert et al., 2003). Since the validation of methods is crucial, a recent comparison of FISH and qPCR as tools to quantify the abundance of β-AOB deserves attention. It revealed that FISH and *amoA*-targeted qPCR yielded consistent results, whereas qPCR of 16S rRNA genes underestimated the abundance of β-AOB (Baptista et al., 2014).

These and many other studies used a well-established set of rRNA-targeted oligonucleotide probes to detect all β-AOB or their sublineages by FISH (Adamczyk et al., 2003; Juretschko et al., 2002, 1998; Mobarry et al., 1996; Wagner et al., 1995). However, the most commonly used probes were designed based on a much more limited set of 16S rRNA sequences from β-AOB than is available now, and they do not cover the entire diversity of β-AOB represented in current databases. Moreover, inconsistent *in situ* hybridization patterns were observed, which suggested that some of these probes hybridize to β-AOB outside the expected target groups (Gruber-Dorninger et al., 2015). An incomplete probe coverage and weak specificity can introduce a significant bias in studies that rely on these probes to detect and quantify β-AOB and to distinguish different β-AOB lineages. Hence, an updated set of β-AOB-specific FISH probes is urgently needed.

Here we introduce new β-AOB-specific rRNA-targeted oligonucleotide probes, which complement and refine the existing probe set and enable the identification, visualization, and quantification of all currently known β-AOB lineages. Following probe design and evaluation, the specificity and applicability of the new probes were tested with activated sludge and biofilm samples from municipal and industrial WWTPs.

## 2. Materials and methods

### 2.1 In silico *design of 16S rRNA-targeted probes and phylogenetic analyses*

The new rRNA-targeted oligonucleotide probes (Table 1) were designed using the “probe design” and “probe match” functions of the ARB software package (version 6.0.6) (Ludwig et al., 2004) and the SILVA Ref_NR99 (release 132) SSU rRNA sequence database (Quast et al., 2013). The database was amended with 46 additional full-length 16S rRNA gene sequences of β-AOB from full-scale WWTPs in Denmark (Dueholm et al., 2020), which were retrieved from the MiDAS database (McIlroy et al., 2015; Nierychlo et al., 2020). The “probe match” tool of ARB was also used to evaluate the target group coverage and specificity of the previously published and the new FISH probes for β-AOB. These analyses, and the probe specificity checks outside of the β-AOB, used SILVA Ref_NR99 (release 138) amended with the 46 sequences from MiDAS (see above). A maximum likelihood phylogenetic tree was calculated, based on an alignment (SINA aligner, Pruesse et al., 2012) of 505 reference sequences of β-AOB from the aforementioned databases, using W-IQ-Tree (Trifinopoulos et al., 2016) with 1,000 bootstrap iterations. A previous phylogenetic study had found that different treeing methods (including maximum likelihood) yielded consistent tree topologies at the level of the major β-AOB lineages and with the 16S rRNA gene as marker (Purkhold et al., 2003). TIM3e+G4 was determined by ModelFinder to be the best fitting base substitution model for the calculation (Kalyaanamoorthy et al., 2017). The resulting tree was visualized with iTOL (Letunic and Bork, 2019).

**Table 1.**
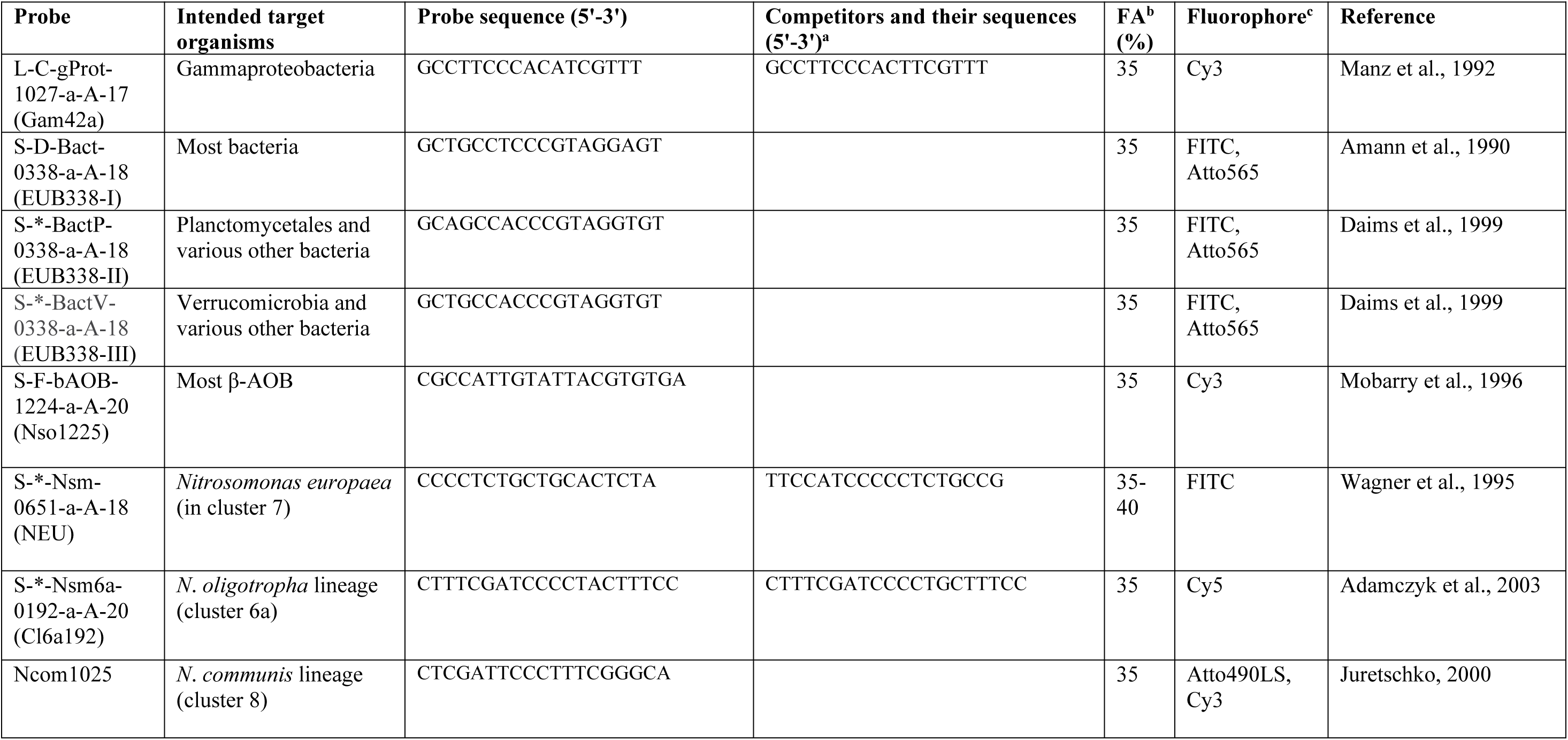
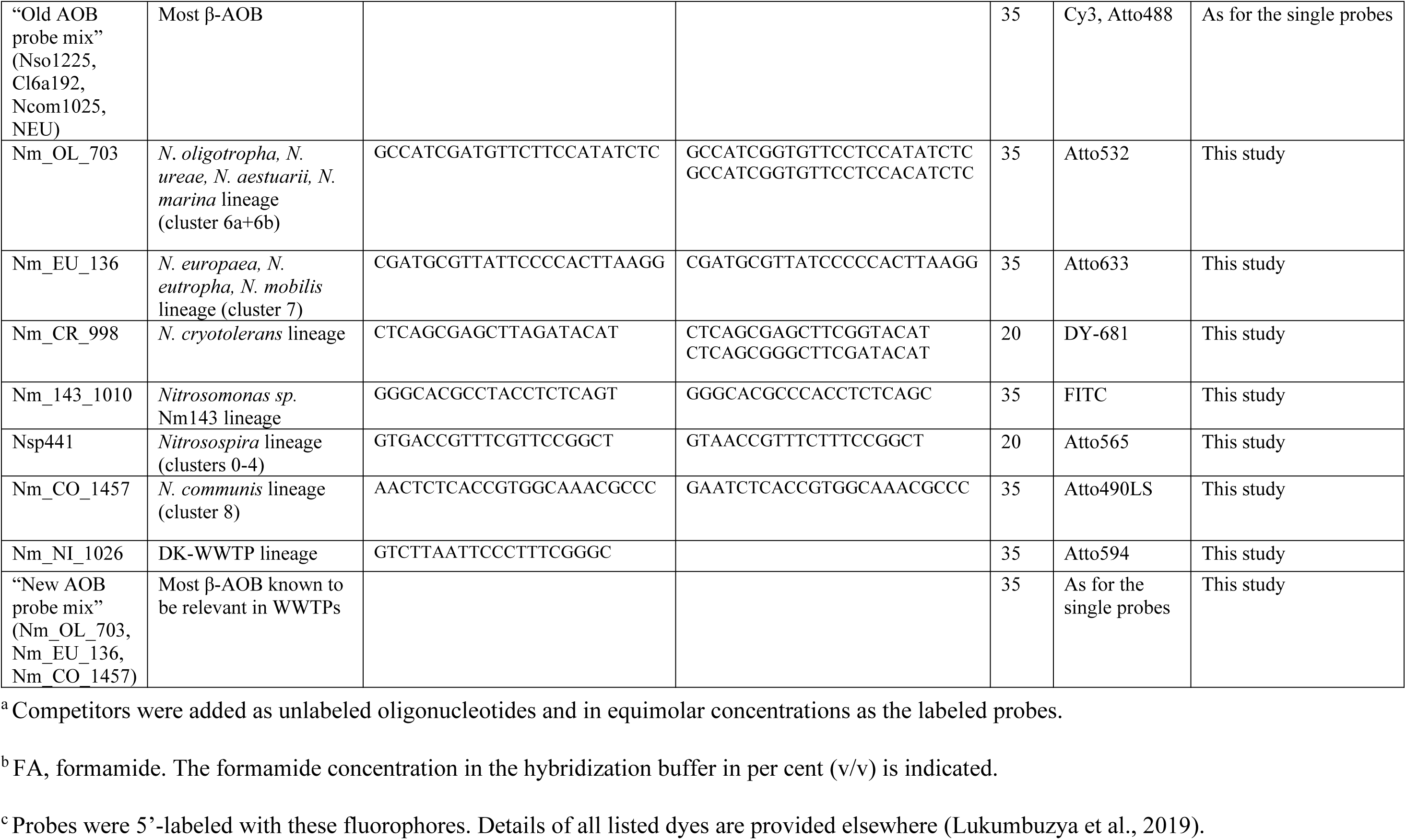
rRNA-targeted oligonucleotide probes used in this study. Probe Gam42a binds to the 23S rRNA, whereas all other listed probes target binding sites on the 16S rRNA.

Bacterial 16S rRNA amplicon sequence data (V1-3 region) from 88 samples from the five Danish wastewater treatment plants (Table 2) were retrieved from the MiDAS database (McIlroy et al., 2015) and used to determine the mean amplicon abundances of β-AOB classified as *Nitrosomonas* and *Nitrosospira*. Amplicon reads were dereplicated and formatted for use in the USEARCH/UNOISE workflow (Edgar, 2016a). Dereplicated reads were used to generate amplicon sequence variants using the USEARCH (version 10) “unoise3” tool with default settings. Taxonomy was assigned using the SINTAX classifier (Edgar, 2016b) in USEARCH (version 11), using the SILVA (release 138) taxonomy. The results were analyzed in R (R Core Team, 2016) using the ampvis2 R package 2.5 (https://github.com/MadsAlbertsen/ampvis2).

**Table 2.** Overview of the WWTPs, which were the source of activated sludge samples analyzed in this study.

| WWTP<br>(abbreviations) | Location<sup>a</sup> | Sampling date<br>(month/year) | Type of WWTP/reactor |
| --- | --- | --- | --- |
| KNB | Klosterneuburg, AT | 10/2014, 10/2015 | Municipal. Sequencing batch reactor that treats reject water from sludge dewatering after anaerobic digestion. |
| SBBR1 | Ingolstadt, DE | 3/1998 | Municipal. Previous pilot-size sequencing biofilm batch reactor that treated reject water from sludge dewatering after anaerobic digestion. |
| ING | Ingolstadt, DE | 3/2011 | Municipal. Denitrifying-nitrifying tank. |
| CP Kelco | Lille Skensved, DK | 12/2014 | Industrial. 300,000 PE full-scale wastewater treatment plant with biological nutrient removal, recirculation and anaerobic digestion. |
| Esbjerg E | Esberg, DK | 8/2016 | Municipal. 150,000 PE full-scale wastewater treatment plant with biological nutrient removal, recirculation, primary settling and anaerobic digestion. |
| Kalundborg | Kalundborg, DK | 8/2016 | Municipal. 50,000 PE full-scale wastewater treatment plant with enhanced biological phosphorus removal, intermittent aeration. |
| Lundtofte | Lyngby, DK | 8/2010 | Municipal. 150,000 PE full-scale wastewater treatment plant with enhanced biological phosphorus removal, intermittent aeration, primary settling, and anaerobic digestion. |
| Odense NW | Odense, DK | 11/2014 | Municipal. 51,000 PE full-scale wastewater treatment plant with biological nutrient removal and intermittent aeration, primary settling and anaerobic digestion. |

<sup>a</sup> Abbreviations: AT, Austria; DE, Germany; DK, Denmark.

### 2.2 Cultivation and fixation of β-AOB pure cultures

Pure cultures of the β-AOB *Nitrosospira briensis* Nsp1, *Nitrosospira multiformis* NI13, *Nitrosomonas europaea* Nm50, *Nitrosomonas eutropha* Nm57, *Nitrosomonas oligotropha* Nm75, and *Nitrosomonas sp.* Nm51 (an unnamed species from the *Nitrosomonas marina* lineage) were grown as described earlier (Koops et al., 1991). *Nitrosomonas communis* Nm2 was grown in a modified AOB medium according to Zhou *et al.* (2019). All cultures were harvested (~40 ml) during the logarithmic growth phase and centrifuged (4,000×g, room temperature, 20 min) to collect the biomass. The supernatant was removed, and the cell pellets were resuspended in a 3% (w/v) formaldehyde solution for fixation (1 h at room temperature) as detailed elsewhere (Daims et al., 2005). The fixed cultures were washed twice in a 1× PBS (phosphate-buffered saline) solution, centrifuged 12,000×g at room temperature for 8 min, and then resuspended in a 1:1 mixture of 1× PBS and 96% (v/v) ethanol, and stored at −20°C until further processing (Daims et al., 2005).

### 2.3 Sampling and fixation of activated sludge

Activated sludge or biofilm (detached, suspended particles) samples taken at several WWTPs in Austria, Germany, and Denmark were analyzed (Table 2). The samples were centrifuged (20,817×g, 4 °C, 15 min), the supernatant was removed, and the sludge was resuspended in a 2% (v/v; samples from SBBR1, KNB, and ING) or 4% (v/v; all other samples) formaldehyde solution for fixation (3 h, 4 °C) (Lukumbuzya et al., 2019; Nielsen, 2009). The sludge samples were subsequently washed twice in 1× PBS, resuspended in a 1:1 mixture of 1× PBS and 96% (v/v) ethanol, and stored at −20°C until further processing.

### 2.4 Recombinant 16S rRNA expression for Clone-FISH

We could not obtain cells of the isolates *Nitrosomonas cryotolerans* (targeted by the new probe Nm_CR_998) and *Nitrosomonas sp.* Nm143 (targeted by probe Nm_143_1010) (Table 1). Furthermore, no isolate is available from a *Nitrosomonas communis*-related cluster detected in some WWTPs in Denmark (Table 2), for which we designed the new probe Nm_NI_1026 (Table 1). In order to evaluate the new probes that are specific for these β-AOB and their close relatives, the respective 16S rRNA was heterologously expressed in *E. coli* for Clone-FISH (Schramm et al., 2002). Briefly, synthetic Strings DNA fragments (ThermoFisher Scientific) of full-length 16S rRNA genes were cloned into *E. coli* NovaBlue competent cells using the Novagen pETBlue-1 Perfectly Blunt Cloning Kit (Merck KGaA). The *E. coli* cells were grown to an OD of 0.3-0.4, 1 mM of IPTG was added to the cultures, and the cells were incubated (1 h, 200 r.p.m., 37 °C). Subsequently, chloramphenicol (170 mg l^-l^) was added to increase the intracellular accumulation of RNA (including the heterologously expressed 16S rRNA) (Schramm et al., 2002) and the cells were incubated at 4 °C for 4 hours. Finally, the cells were fixed in formaldehyde as described above for the β-AOB isolates.

### 2.5 Fluorescence *in situ* hybridization, microscopy, and digital image analysis

FISH of all β-AOB pure cultures, *E. coli* cells containing recombinant 16S rRNA (Clone-FISH), and activated sludge samples was performed according to the standard protocol for FISH with rRNA-targeted oligonucleotide probes (Table 1) (Daims et al., 2005; Manz et al., 1992). Briefly, probe solutions had a concentration of 5 pmol µl^-1^ and were applied at a ratio of 1:10 (v/v) in hybridizaton buffer. If applicable, unlabeled competitor oligonucleotides (Table 1) were used in equimolar concentrations as the probes. Hybridizations were performed at 46 °C for 2 hours. After hybridization, samples were washed in washing buffer for 10 min at 48 °C and shortly dipped into ice-cold MilliQ water. All hybridized samples were also stained with DAPI (4′,6-diamidino-2-phenylindole). For this purpose, 10 µl of 10 mg/ml DAPI was spotted onto hybridized samples, incubated for 5 min at room temperature, and subsequently washed away by dipping samples in 96% (v/v) ethanol. Samples were analysed immediately or stored at −20°C.

Fluorescence micrographs of probe-labelled cells were acquired using an inverted Leica TCS SP8X confocal laser scanning microscope (CLSM). The CLSM was equipped with a UV 405 diode and a supercontinuum white light laser, two photomultiplier (PMT) detectors, three hybrid (HyD) detectors, and the Leica Application Suite AF 3.2.1.9702 or Leica Application Suite X 3.5.6.21594. The settings for excitation and emission wavelengths were adjusted to match the respective fluorochromes (Table 1) as described elsewhere for multicolor FISH (Lukumbuzya et al., 2019). The digital image analysis and visualization software *daime* (version 2.2) (Daims et al., 2006) was used to project 3D confocal *z*-stacks.

To evaluate probe dissociation profiles, β-AOB pure cultures or *E. coli* cells (for Clone-FISH) were hybridized to the respective probes with increasing concentrations of formamide [0 to 70% (v/v)] in the hybridization buffer and corresponding salt concentrations in the wash buffer (Manz et al., 1992). If applicable, competitor oligonucleotides (Table 1) were included. Images for inferring probe dissociation profiles were recorded using the same CLSM settings for all parameters (laser power, confocal pinhole size, and “smart gain”). The probe dissociation profiles were determined, based on the mean fluorescence intensities of the probe-labelled cells, by using the respective tool of the *daime* software. The data were plotted in R, and approximated probe dissociation curves were obtained by non-linear regression with a sigmoidal model.

For the quantification of relative biovolume fractions, activated sludge samples were hybridized to a β-AOB specific probe mix and to the EUB338 I-III probe mix (Table 1), both labeled with different fluorochromes (Daims and Wagner, 2007). The two β-AOB-specific probe mixtures used consisted of previously published or newly designed probes, respectively (Table 1). In the “old” β-AOB probe mix, all probes were labelled with Cy3, and their fluorescence signals were recorded together in the same image. In the “new” β-AOB probe mix, the probes were labelled individually with different dyes, and the recorded images of these fluorescence signals were merged *in silico* for quantifying the relative biovolume fractions of β-AOB. Ten to 40 pairs of images containing the specific probe or EUB338 I-III probe signals, respectively, were acquired at random positions in a sample (Daims and Wagner, 2007). The CLSM settings were adjusted so that cells in the specific probe images had the same size as their counterparts in the EUB338 I-III images (Daims et al., 2005). The respective tool of the *daime* software was used to measure the biovolume fractions of β-AOB based on these image pairs.

## 3. Results and discussion

### 3.1 Evaluation of existing FISH probes targeting β-AOB

Most previous studies using FISH to detect β-AOB *in situ* relied on a set of 16S rRNA-targeted oligonucleotide probes, which were designed as long as 16 to 25 years ago. This set consists of the probe Nsm156 (for various *Nitrosomonas* spp.), NEU (for halophilic and halotolerant *Nitrosomonas* spp.), Cl6a192 (for the *Nitrosomonas oligotropha* lineage), NmV (for *N. mobilis*), Ncom1025 (for *Nitrosomonas communis*), Nsv443 (for the *Nitrosospira* lineage), and the two broad-range probes Nso190 and Nso1225 (for most β-AOB) (Adamczyk et al., 2003; Juretschko, 2000; Juretschko et al., 1998; Mobarry et al., 1996; Wagner et al., 1995). In the present study, we matched the sequences of these probes against a recent 16S rRNA gene sequence database (see 2.1), which contained 505 non-redundant, full-length sequences from cultured and uncultured β-AOB. This *in silico* analysis revealed considerable gaps in the target group coverage for some of the probes, whereas others still showed a surprisingly good coverage (Table 3). In particular, the broad-range probe Nso1225 still covers the vast majority of β-AOB, whereas probe Nso190 (originally also designed to target all β-AOB) has a highly incomplete coverage according to current databases. Probe Nsm156, which should target the genus *Nitrosomonas* (Mobarry et al., 1996), still covers a large fraction of this genus. Probe Cl6a192 for the *N. oligotropha* lineage (cluster 6a) covers only ~50% of its target group (Table 3). Furthermore, unexpected hybridization patterns had previously been observed for the probes Cl6a192, NEU, and Ncom1025: although these probes target different lineages of β-AOB, a large proportion of their signals overlapped in FISH experiments with activated sludge (Gruber-Dorninger et al., 2015). These observations are consistent with a lack of specificity of probes NEU and Ncom1025, which becomes apparent when non-target β-AOB-sequences with 1-2 weak nucleotide mismatches to the probes are taken into account (Table 3). Such weak mismatches are often difficult to discriminate in FISH without competitor oligonucleotides. In this case, probe NEU potentially covers ~30% of cluster 6a (the target group of probe Cl6a192) and would probably also bind to members of the *N. mobilis* lineage within cluster 7. Notably, this unspecific hybridization of probe NEU would not be prevented by the published competitor (Table 1). Probe Ncom1025 would also potentially detect the majority of the *N. mobilis* lineage and could thus bind to the same organisms as probe NEU (Table 3). In our study, unspecific probe hybridization was confirmed in tests using biofilm and activated sludge samples from two bioreactors, a sequencing batch biofilm reactor (SBBR1) and the WWTP of Klosterneuburg, Austria (KNB) (Fig. 1A, C; Table 2). Moreover, in one additional sludge from an industrial WWTP (CP Kelco, Table 2), the probe mix consisting of previously published AOB probes (Table 1) detected numerous microbial cells that were arranged as tetrads within loose aggregates (Fig. 2A). The morphology of these organisms was very dissimilar from the usual size and shape of the β-AOB cells and cell clusters, which were also present in this sludge (Fig. 2A). Moreover, a test hybridization revealed that the tetrad-shaped cells were also labelled by probe Gam42a (Fig. S1), suggesting that these organisms were Gammaproteobacteria and unspecifically labelled by the previously published AOB probes. In summary, the *in silico* evaluation and test hybridizations demonstrated that several of the previously published FISH probes for β-AOB suffer from an insufficient target group coverage and specificity. As a consequence, further use of the affected probes for the *in situ* identification and quantification of the respective β-AOB groups should be performed with caution by taking into account their actual specificities (Table 3).

**Table 3.**
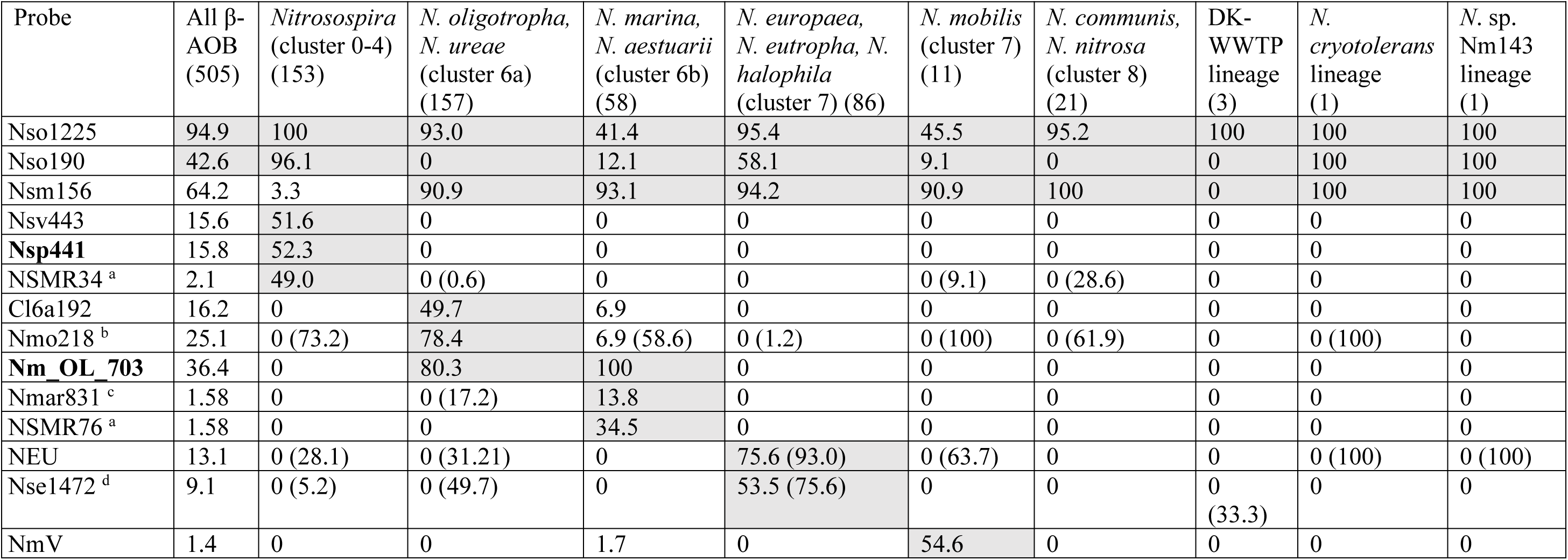
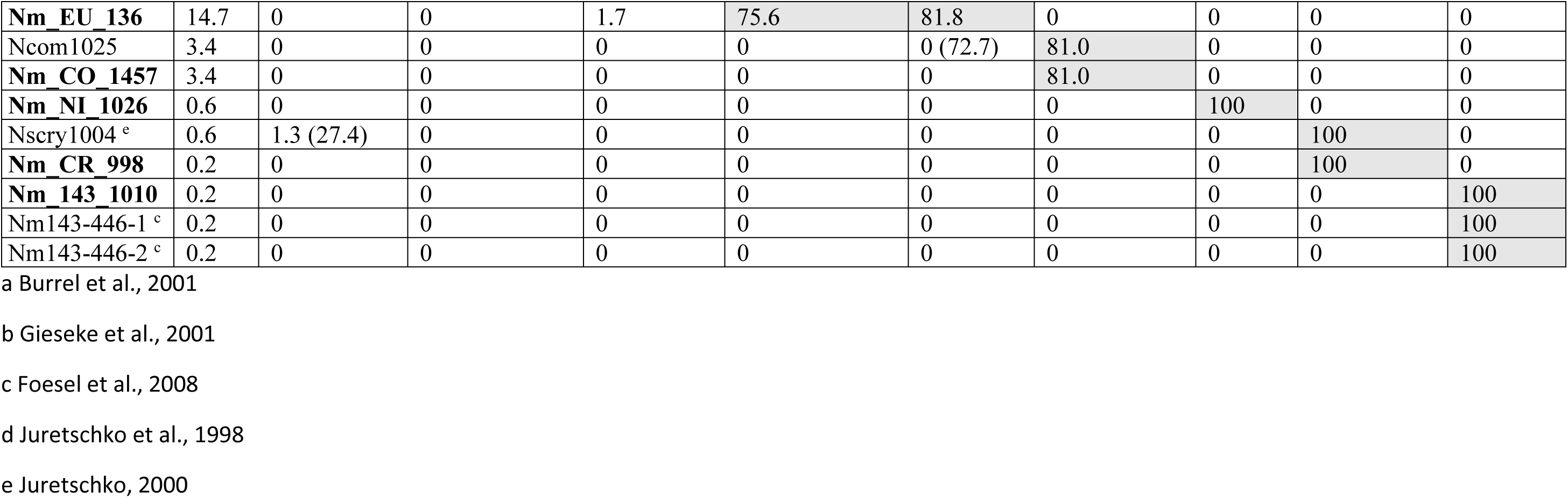
*In silico* analysis of the target group coverage and specificity of previously published and newly designed 16S rRNA-targeted probes for β-AOB. The names of new probes are printed in bold. Numbers in the table header (in parentheses) are the numbers of analyzed full-length 16S rRNA gene sequences from the respective lineage in the SILVA Ref_NR99 (release 138) and MiDAS databases. Numbers in all other rows indicate the fractions (in per cent) of the lineages that are targeted by the probes without any nucleotide mismatch. Numbers in parentheses indicate the fractions (in per cent) targeted by the probes with up to 1.5 weighted mismatches according to the ARB “probe match” tool. Table cells of the intended probe target groups are marked grey. Those non-target organisms of a probe, which are discriminated by the competitor listed in Table 1, are not included in the listed fractions.

**Figure 1.**
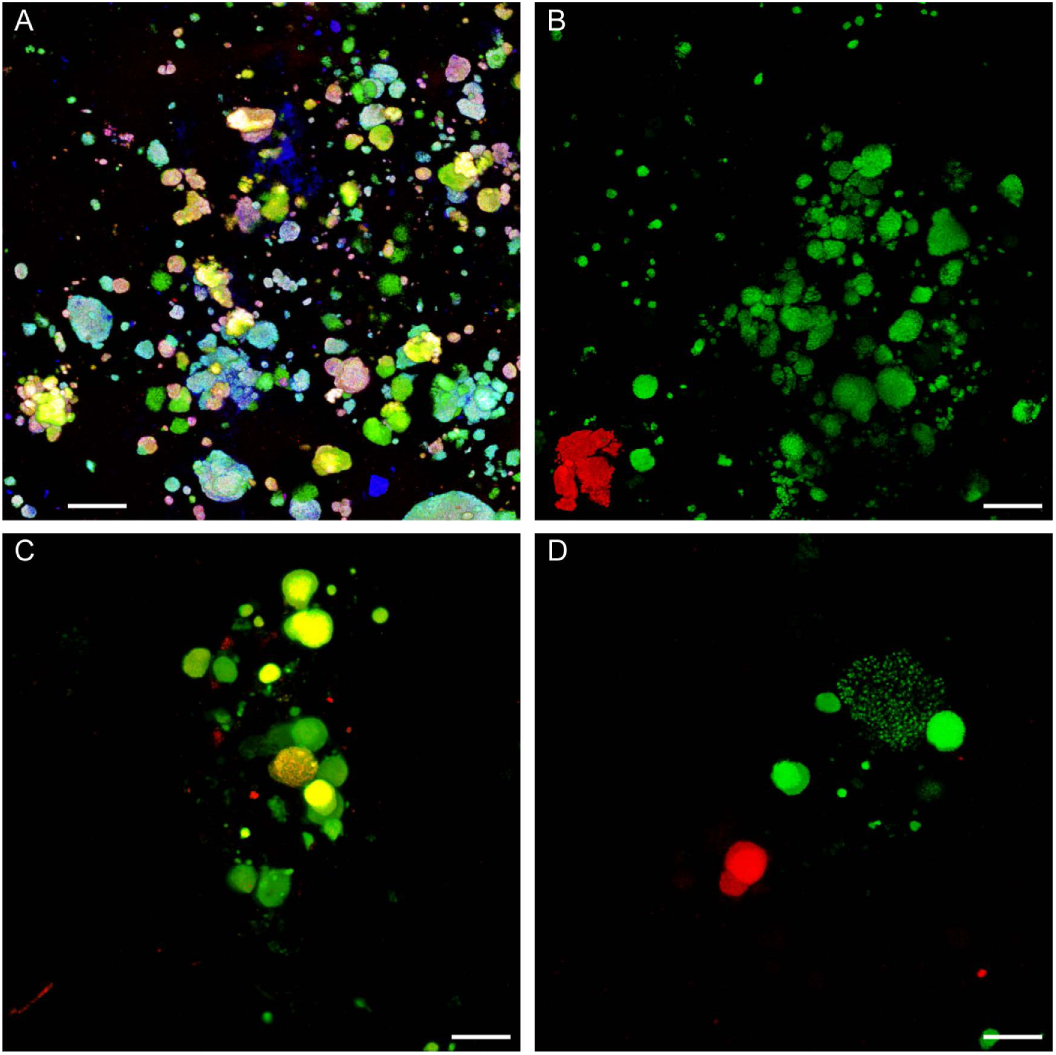
Comparison of hybridization patterns of selected previously published and newly designed FISH probes targeting β-AOB. (**A**) FISH of a suspended biofilm particle from reactor SBBR1 with probes NEU (red), Cl6a192 (green), and Ncom1025 (blue). Mixed colors (cyan, magenta, yellow, white) indicate binding of multiple probes to the same microcolonies of β-AOB. (**B**) Sample of the same biofilm as in panel A after FISH with probes Nm_EU_136 (red) and Nm_OL_703 (green). No cross-hybridization was observed. Probe Nm_CO_1457 targeting the same lineage as probe Ncom1025 was also applied, but no signals were recorded, indicating that probe Ncom1025 (panel A) detected non-target β-AOB in this sample. (**C**) FISH of sludge from WWTP KNB with probes NEU (red) and Cl6a192 (green). Yellow indicates binding of both probes to the same microcolonies of β-AOB. (**D**) Sample of the same sludge as in panel C after FISH with probes Nm_EU_136 (red) and Nm_OL_703 (green). No cross-hybridization was observed. (**A-D**) All panels show projections of 3D confocal *z*-stacks. Bar = 20 µm.

**Figure 2.**
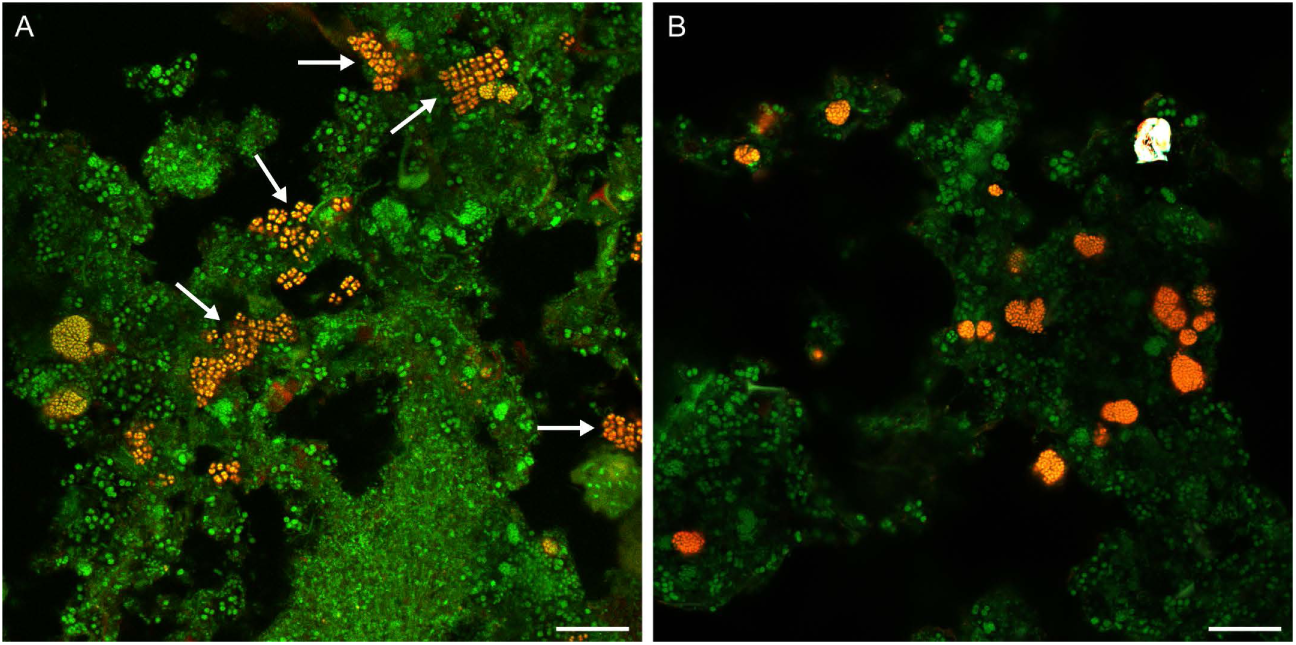
Detection of β-AOB in sludge from the WWTP CP Kelco using previously published and newly designed AOB probe mixtures. (**A**) Application of the old AOB probe mix (red) and probes EUB338 I-III (green). Orange indicates binding of the AOB- and EUB338-probes to the same cells. Note the detection of numerous tetrad-shaped cells (arrows) by the old AOB mix and the EUB338 probes. (**B**) Application of the new AOB probe mix (red) and probes EUB338 I-III (green). The white object in the upper right corner is an artifact. (**A, B**) Bar = 20 µm. Details of the probe mixtures are provided in Table 1.

### 3.2 Design of a refined, β-AOB cluster-specific 16S rRNA-targeted oligonucleotide probe set

Based on 16S rRNA and *amoA* gene phylogenies, a subdivision of the β-AOB into distinct lineages (“clusters”) was suggested previously (Koops et al., 2003; Purkhold et al., 2003, 2000; Stephen et al., 1996). In that phylogenetic framework, members of the genus *Nitrosospira* belong to the clusters 0 to 4, whereas the remaining clusters 5 to 8 are formed by different lineages of the genus *Nitrosomonas* (Purkhold et al., 2000). Two additional, separate lineages were formed by *Nitrosomonas cryotolerans* and the estuary isolate *Nitrosomonas* sp. Nm143 (Purkhold et al., 2003, 2000). Notably, the obtained tree topologies did not support a clear separation of the *Nitrosospira* and *Nitrosomonas* genera; instead, the analyses indicated that the currently defined genus *Nitrosomonas* is not monophyletic within the β-AOB (Purkhold et al., 2003, 2000). Since the individual *Nitrosomonas* clusters nevertheless represent stable lineages in bootstrap analyses (Purkhold et al., 2003), and a thorough phylogenetic and taxonomic reevaluation of the β-AOB is still pending, we retain the established nomenclature in this study.

Using the β-AOB clusters as a framework we designed six new, 16S rRNA-targeted oligonucleotide probes that are specific for *Nitrosomonas* cluster 6 (including both 6a with *N. oligotropha* and *N. ureae*, and 6b with *N. marina* and *N. aestuarii*), cluster 7 (with *N. europaea*, *N. eutropha*, and *N. mobilis*), cluster 8 (with *N. communis* and *N. nitrosa*), the *N. cryotolerans* cluster, the *N*. sp. Nm143 cluster, and a new environmental cluster “DK-WWTP” related to cluster 8 (Fig. 3). In addition, the old probe Nsv443, which offered good coverage of the target clade but unsatisfactory target specificity, was replaced with the new probe Nsp441 for the genus *Nitrosospira* (comprising clusters 0-4) (Fig. 3). For most of the new probes, we designed competitor oligonucleotides that help discriminate non-target organisms which possess only one or few weak base mismatches to the probe sequence in their 16S rRNA. These competitors were used as unlabeled oligonucleotides and in equimolar concentrations as the labeled probes, in FISH experiments. Details of the probes, their target groups, and the competitors are listed in Table 1. According to an *in silico* analysis, the new probes in combination with the designated competitors (Table 1) display a very good coverage and specificity for their target β-AOB clusters, including cultured isolates as well as environmental sequences from uncultured β-AOB (Table 3). Notably, the newly designed probes (with competitors) also display a very high specificity with respect to non-target matches outside of the β-AOB (*in silico* evaluation based on the SILVA SSU NR release 138 and the ARB “probe match” tool with 0-2 weighted mismatches as search criterion). Merely for probe Nm_143_1010, less than ten non-target betaproteobacterial sequences were found that are not covered by the competitor for this probe (Table 1), and probe Nsp441 (Table 1) might hybridize with less than 60 non-target betaproteobacteria (mainly from the genera *Hydrogenophaga* and *Gallionella*). However, the number for Nsp441 is low compared to the previously published probe Nsv443 that would potentially detect more than 500 non-target organisms. Considering that any environmental sample likely contains non-target organisms, which are not present in the current sequence databases, we recommend to use the newly designed β-AOB cluster-specific probes in combination with the broad-range probe Nso1225 (Table 3) labelled with a different fluorochrome. Cells detected by both probes should represent the targeted β-AOB lineage. In order to identify the optimal hybridization stringency for each probe, the dissociation profiles of the probes were determined by performing FISH at increasing hybridization and washing stringencies (Manz et al., 1992). Where possible, pure culture cells of the target β-AOB were used in these experiments. As isolates of *N. cryotolerans*, *N*. sp. Nm143, and the uncultured cluster DK-WWTP were not available, their 16S rRNA genes were heterologously expressed in *E. coli* (Schramm et al., 2002) to determine the dissociation profiles of the respective probes. For all probes, we obtained sigmoid dissociation profiles that were suitable to identify the highest stringency, which still yields bright fluorescence signals with the target organisms (Fig. S2 and S3, Table 1). Probes that need different hybridization stringencies can be used in the same experiment by performing the most stringent hybridization and washing steps first and then the other steps in order of decreasing stringency (e.g., Daims et al., 2005).

**Figure 3.**
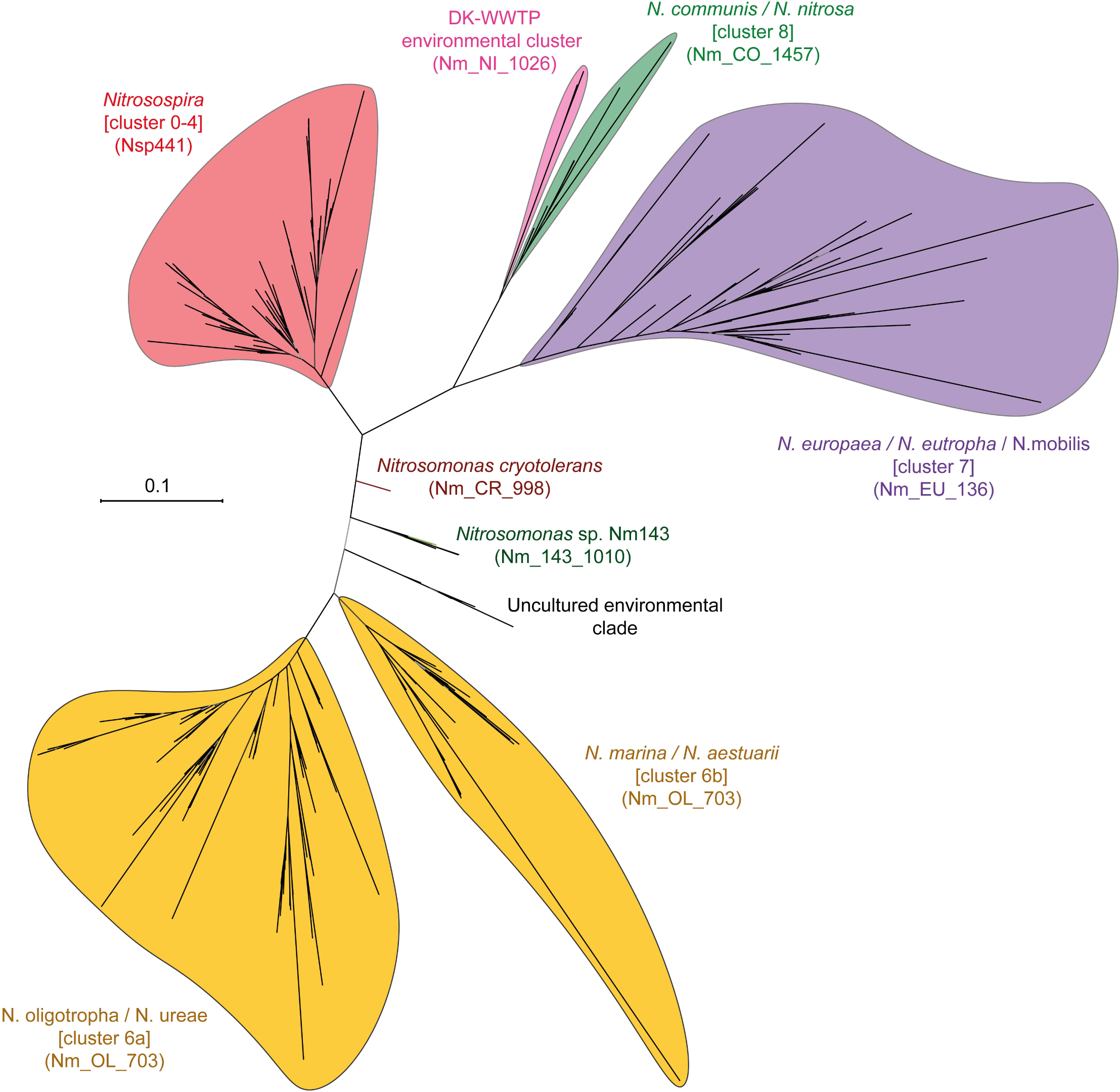
Unrooted maximum likelihood tree showing the major β-AOB lineages. Cluster designations, according to Purkhold *et al*. (2000), are indicated in brackets. The names of newly designed 16S rRNA-targeted oligonucleotide probes are indicated in parentheses. The scale bar depicts 0.1 estimated substitutions per nucleotide.

All probes were also hybridized to non-target β-AOB cells (or the respective recombinant *E. coli* cells) to test for unspecific hybridization. In these experiments, no fluorescence signal was observed for any probe at the optimal stringency and in the presence of the competitor oligonucleotides (Fig. S4).

### 3.3 Detection and quantification of β-AOB in activated sludge

The applicability of the new probe set was tested in FISH experiments with nitrifying activated sludge samples from different WWTPs in Austria, Germany, and Denmark (Table 2). In particular, the newly designed probes targeting *Nitrosomonas* cluster 6 (Nm_OL_703) and cluster 7 (Nm_EU_136) were applied to the same sludge samples that we had already used to confirm the cross-hybridization of the previously published probes Cl6a192, NEU, and Ncom1025 (Fig. 1). With the two new probes, no overlapping fluorescence signals were observed, and the hybridization patterns appeared to be completely consistent (Fig. 1B, D). This result is in agreement with the *in silico* analysis, which predicted that probes Nm_OL_703 and Nm_EU_136 do not target the same sequences in the database (Table 3). Furthermore, when the “new AOB probe mix” (Table 1) was used to detect β-AOB in the sludge from CP Kelco, the conspicuous tetrad-shaped cells, detected by the “old AOB probe mix” were not labelled anymore. Instead, the “new probe mix” detected exclusively cells and cell clusters that displayed the typical morphology of β-AOB, which has been observed in numerous studies of nitrifiers in WWTPs and isolated β-AOB (Daims et al., 2001; e.g., Juretschko et al., 1998; Koops and Pommerening-Röser, 2001) (Fig. 2B). The new probe Nm_NI_1026, which targets the novel uncultured lineage DK-WWTP (Fig. 3), was applied to activated sludge from WWTP Esbjerg East where it showed specific signals with a morphology similar to that typically portrayed by β-AOB (Fig. 4). We did not have access to environmental samples that were suitable for FISH and contained the target β-AOB of the new probes Nm_CR_998, Nm_143_1010, Nm_CO_1457, and Nsp441. However, these probes were successfully evaluated using pure β-AOB cultures or Clone-FISH. In addition, they did not unspecifically detect other bacterial populations in activated sludge or biofilm (Fig. 1 for Nm_CO_1457; Fig. S5 for Nm_CR_998, Nm_143_1010, and Nsp441). Under the assumption that those probes, which could only be tested by Clone-FISH, will also bind to the native ribosomes of their target organisms, all probes should be suitable for FISH analyses of environmental AOB communities.

**Figure 4.**
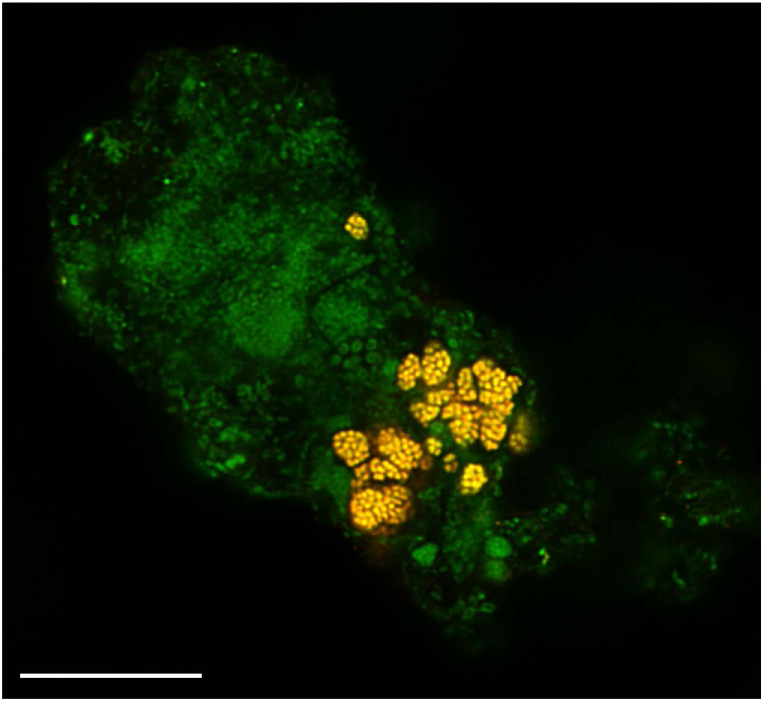
Detection of the novel *Nitrosomonas* lineage DK-WWTP in activated sludge from the WWTP Esbjerg East. Applied probes were EUB338 I-III (green) and NM_NI_1026 (red). Yellow indicates binding of NM_NI_1026 and the EUB338-probes to the same cells. Bar = 20 µm.

A quantitative comparison of the “old” and “new” AOB probe mixes (Table 1) was carried out with activated sludge samples from seven WWTPs (Table 4). For six of these sludge samples, highly similar biovolume fractions of β-AOB were measured by quantitative FISH with either probe set (Table 4). Only for the CP Kelco sludge we obtained a much lower biovolume fraction of β-AOB with the newly designed probe set than with the previously published probes (Table 4). This difference is most likely explained by the better specificity of the newly designed probes, which did not stain the abundant tetrad-shaped cells in this sludge (Fig. 2). For all samples and experiments, high “congruency” values >90% were obtained (Table 4). This value indicates that the fluorescence signals in the β-AOB probe mix images occupied almost exactly the same area as their counterparts in the general bacterial probe (EUB338 I-III) images. Thus, the β-AOB probes did not detect large amounts of non-bacterial cells and did not bind excessively to non-microbial particles in the samples. For the five Danish WWTPs, the results of quantitative FISH gained independent support from similarly low relative abundances of β-AOB in bacterial 16S rRNA gene amplicon datasets (Table 4). Our results confirm that the new β-AOB probe mix (Table 1) is suitable to detect and quantify β-AOB in WWTPs, and, importantly, the data for CP Kelco show that the new mix also offers a better specificity for β-AOB than the previously published probes (Fig. 2, Fig. S1).

**Table 4.**
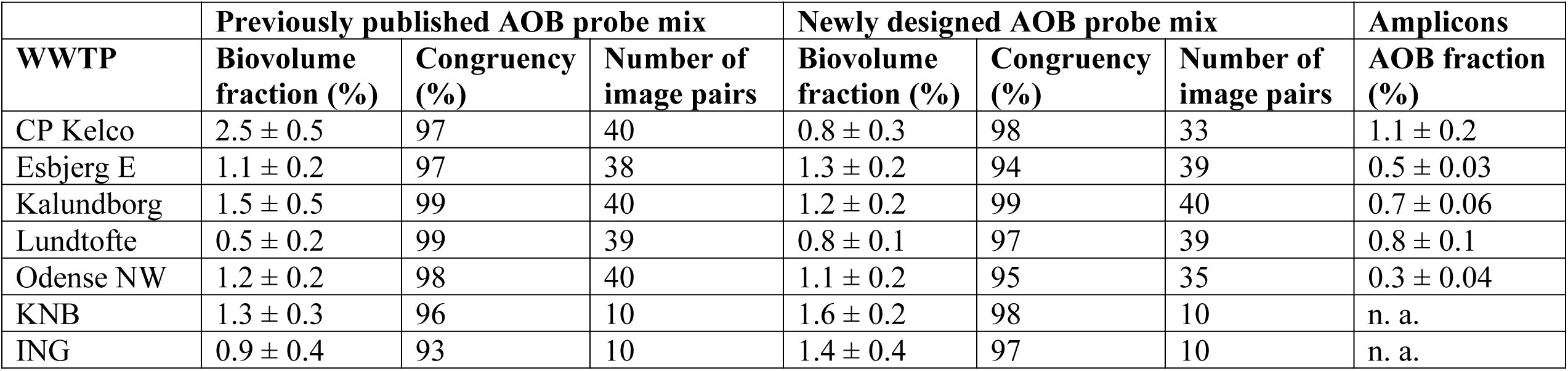
Quantification of the biovolume fractions of β-AOB in seven WWTPs using the old and the new β-AOB probe mixtures. The error values are the standard error of the mean biovolume fraction in the indicated number of image pairs. The mean relative abundances of β-AOB in bacterial 16S rRNA gene amplicon datasets from the Danish WWTPs, as retrieved from the MiDAS database, are listed for comparison (error values are the standard error of the mean). Abbreviation: n. a., not available.

| WWTP | Previously published AOB probe mix |  |  | Newly designed AOB probe mix |  |  | Amplicons |
| --- | --- | --- | --- | --- | --- | --- | --- |
|  | Biovolume fraction (%) | Congruency (%) | Number of image pairs | Biovolume fraction (%) | Congruency (%) | Number of image pairs | AOB fraction (%) |
| CP Kelco | $2.5 \pm 0.5$ | 97 | 40 | $0.8 \pm 0.3$ | 98 | 33 | $1.1 \pm 0.2$ |
| Esbjerg E | $1.1 \pm 0.2$ | 97 | 38 | $1.3 \pm 0.2$ | 94 | 39 | $0.5 \pm 0.03$ |
| Kalundborg | $1.5 \pm 0.5$ | 99 | 40 | $1.2 \pm 0.2$ | 99 | 40 | $0.7 \pm 0.06$ |
| Lundtofte | $0.5 \pm 0.2$ | 99 | 39 | $0.8 \pm 0.1$ | 97 | 39 | $0.8 \pm 0.1$ |
| Odense NW | $1.2 \pm 0.2$ | 98 | 40 | $1.1 \pm 0.2$ | 95 | 35 | $0.3 \pm 0.04$ |
| KNB | $1.3 \pm 0.3$ | 96 | 10 | $1.6 \pm 0.2$ | 98 | 10 | n. a. |
| ING | $0.9 \pm 0.4$ | 93 | 10 | $1.4 \pm 0.4$ | 97 | 10 | n. a. |

## 4. Conclusion

The newly designed 16S rRNA-targeted oligonucleotide probes offer a similar or better coverage of different β-AOB lineages than the established probes (Table 3), and they do not show the unspecific hybridization patterns that were observed for some of the previously designed probes when applied to complex microbial communities (Fig. 1 and 2). However, they are not designed to completely replace the established FISH probes for β-AOB. For example, probes Nso1225 and Nsm156 still offer an excellent coverage of their target groups (Table 3), which is why no new broad-range probe for β-AOB was designed in this study. Instead, the new probes can be applied in combination with selected previously published probes to achieve the currently best possible total coverage of β-AOB and to identify specific β-AOB lineages with a high confidence by FISH. Hence, the new probe set will facilitate future studies of β-AOB community composition and population dynamics. It will also enable specific analyses of the spatial localization of members of the different β-AOB lineages in flocs and biofilms. Such spatial analyses can reveal potential niche differentiation and symbiotic interactions with other microorganisms, but were previously hampered by unspecific hybridization patterns of some of the old probes. Although we used only activated sludge and biofilm samples from WWTPs to test the probes in this study, the new probe set can likely also be applied to other types of samples that are suitable for rRNA-targeted FISH. In summary, the new probes will improve virtually all spatial and functional analyses of β-AOB that use FISH to gain insights which can only be obtained by *in situ* approaches.

## Acknowledgements

This research was supported by the Austrian Science Fund (FWF) via project P27319-B21 to H.D. and W1257 to H.D. and M.W., the Comammox Research Platform of the University of Vienna, the Max Planck Society, and the Danish Research Foundation (grant 6111-00617A to P.H.N.).

