## Supplementary Figures S1-S5 for "A refined set of rRNA-targeted oligonucleotide probes for *in situ* detection and quantification of ammonia-oxidizing bacteria"

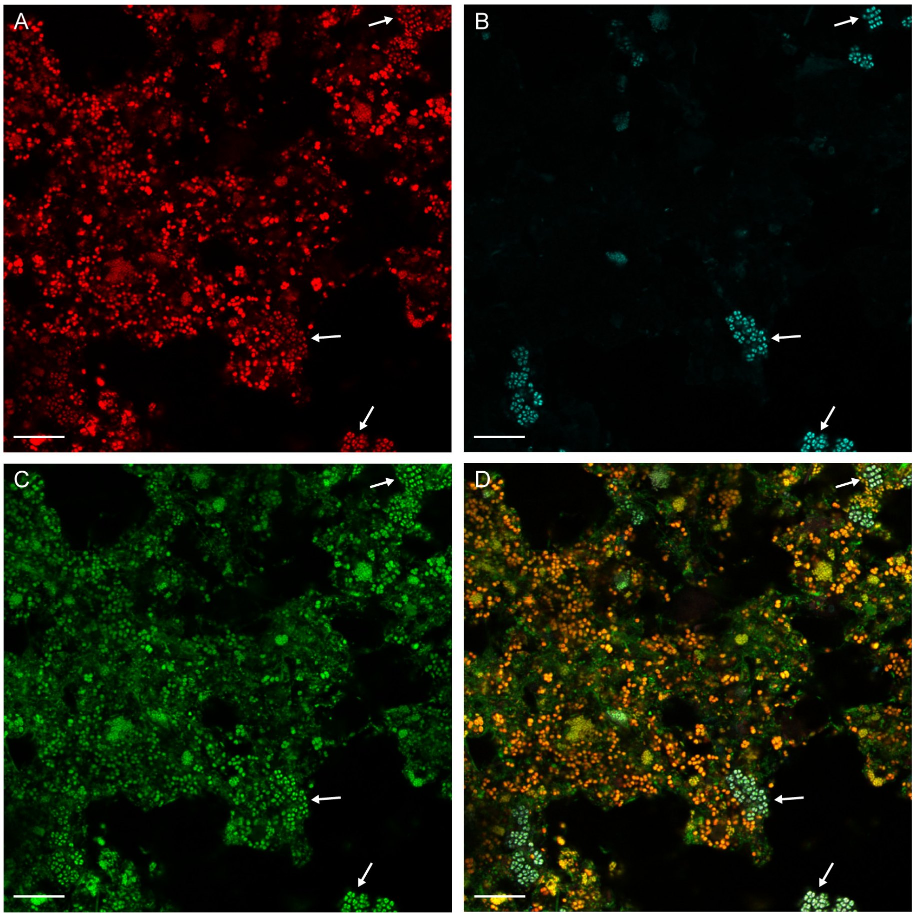


**Figure S1.** Detection of the tetrad-shaped cells in sludge from the WWTP CP Kelco by probe Gam42a and by the previously published AOB probe mixture. (**A**) Application of probe Gam42a (red). (**B**) Application of the previously published AOB mix (cyan). (**C**) Application of probes EUB338 I-III (green). (**D**) Composite image showing all probe signals. Note the binding of probe Gam42a and the old AOB mix to the same tetrad-shaped cells (examples are marked by arrows). Bar = 20 µm. Details of the probes are listed in Table 1 in the main text.

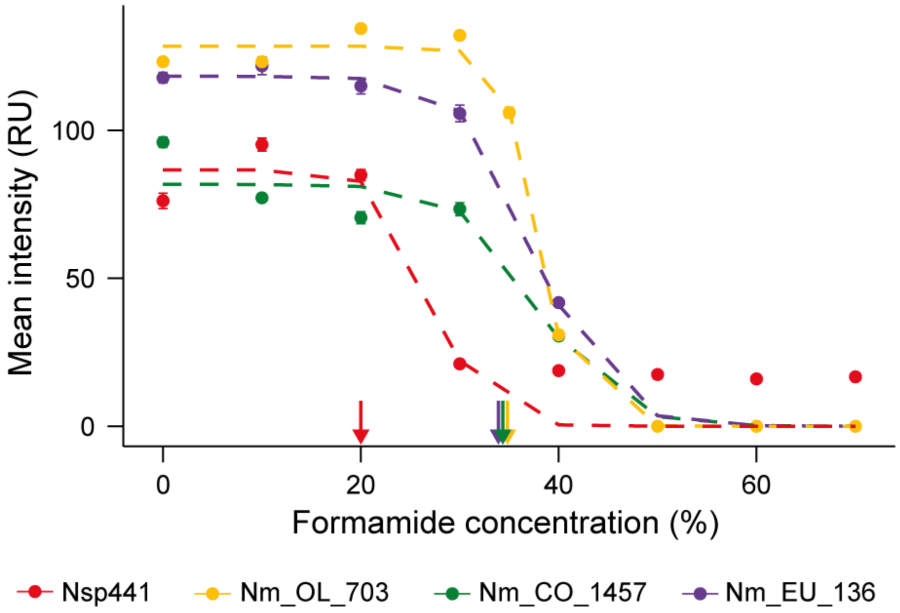


**Figure S2.** Probe dissociation profiles for the newly designed probes Nsp441, Nm_OL_703, Nm_CO_1457, and Nm_EU_136 under increasingly stringent hybridization and washing conditions. These four probes were evaluated using pure culture cells of their target organisms. For each data point, the mean fluorescence intensity of at least 100 cells was measured. Error bars (s.e.m.) are not shown if smaller than the symbols. Dashed lines are regression curves based on a sigmoidal curve fit model. Colored arrows indicate the recommended formamide concentration for the respective probe. Where possible, a formamide concentration of 35% (v/v) was chosen to facilitate combinations of different newly designed and previously published probes in the same FISH experiment. Please note that the values on the *y*-axis cannot be compared between probes, because the analyzed cells most likely differed in ribosome content. For details of the probes please refer to Table 1 in the main text.

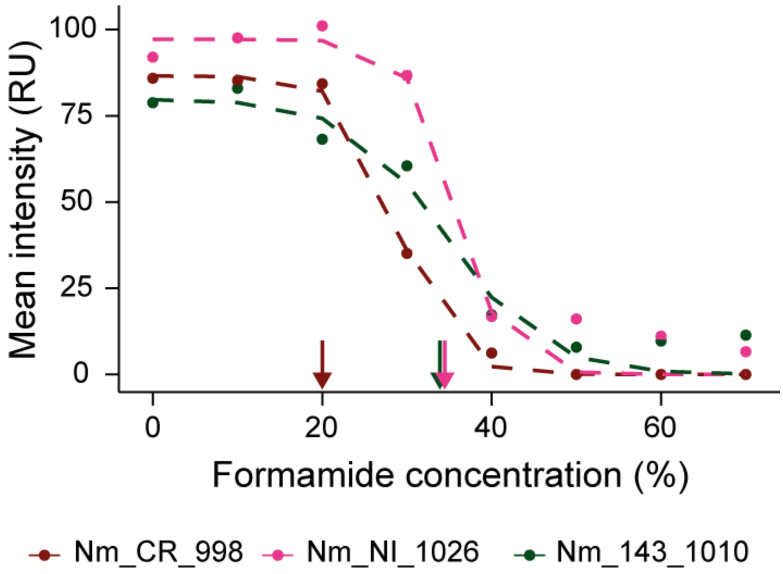


**Figure S3.** Probe dissociation profiles for the newly designed probes Nm_CR_998, Nm_NI_1026, and Nm_143_1010 under increasingly stringent hybridization and washing conditions. These three probes were evaluated using recombinant *E. coli* cells that expressed the target 16S rRNA. For each data point, the mean fluorescence intensity of at least 100 cells was measured. Error bars (s.e.m.) are not shown if smaller than the symbols. Dashed lines are regression curves based on a sigmoidal curve fit model. Colored arrows indicate the recommended formamide concentration for the respective probe. Where possible, a formamide concentration of 35% (v/v) was chosen to facilitate combinations of different newly designed and previously published probes in the same FISH experiment. Please note that the values on the *y*-axis cannot be compared between probes, because the analyzed cells most likely differed in ribosome content. For details of the probes please refer to Table 1 in the main text.

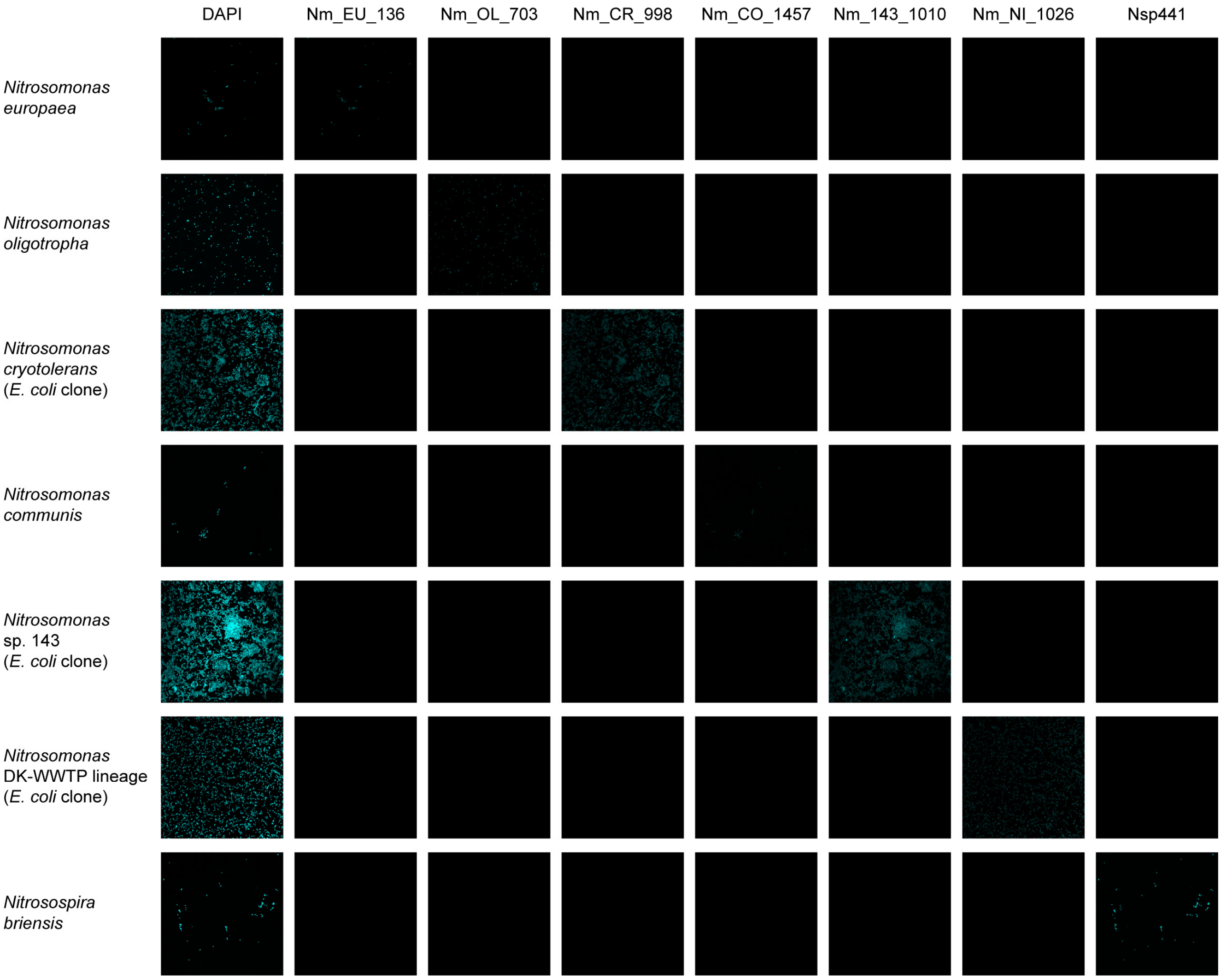


**Figure S4.** Cross-hybridization test with all newly designed FISH probes and the respective target and non-target β-AOB (or recombinant *E. coli* cells). For this test, cells of each organism or *E. coli* clone were simultaneously hybridized to all new probes (each probe labeled with a different fluorochrome) and also stained with DAPI. Sequential hybridizations with 35% and 20% formamide were performed, because probes Nm_CR_998 and Nsp441 require 20% formamide (Table 1 in the main text). Organisms are arranged in rows, probes in columns. Each row shows the same microscopic field of view. The leftmost column shows the cells stained with DAPI. The probes were labeled with the following fluorochromes: Nm_EU_136, Atto633; Nm_OL_703, Atto532; Nm_CR_998, DY-681; Nm_CO_1457, Atto490LS; Nm_143_1010, FITC; Nm_NI_1026, Atto594; Nsp441, Atto565. All fluorescent dyes are false-colored in cyan.

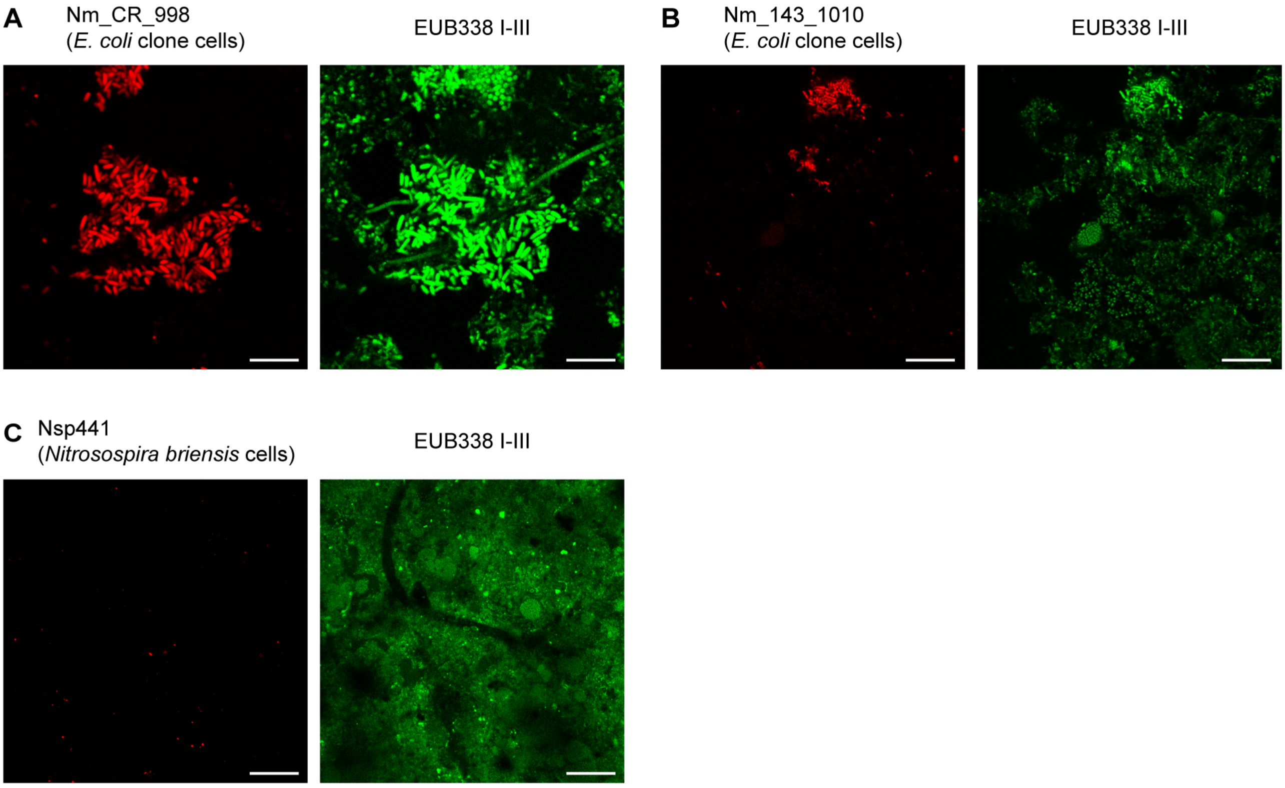


**Figure S5.** Test for unspecific hybridization with activated sludge from WWTP KNB and those three of the new probes, which were not hybridized to activated sludge in the other experiments of this study. Since the KNB sludge did not contain the target β-AOB of these probes, cells from a pure culture of target β-AOB (or recombinant *E. coli* cells) were separately spiked into aliquots of the activated sludge. The spiked samples were simultaneously hybridized to the respective β-AOB-targeted probe (left image, red) and to the EUB338 I-III probe mix (right image, green). As expected, the three probes hybridized to the respective spiked target cells. Unspecific hybridization to other bacterial populations in the sludge was not observed. (**A**) Probe Nm_CR_998 targeting the *Nitrosomonas cryotolerans* lineage; (**B**) probe Nm_143_1010 targeting the *Nitrosomonas* sp. Nm143 lineage; (**C**) probe Nsp441 targeting the *Nitrosospira* lineage. Scale bars = 20 µm.
